# HIF1α inhibition by dual targeting of CDK4/6 and HSP90 reduces cancer cell viability including Rb-deficient cells

**DOI:** 10.1101/2020.10.20.347633

**Authors:** Shuai Zhao, Lanlan Zhou, David T. Dicker, Avital Lev, Shengliang Zhang, Wafik S. El-Deiry

## Abstract

Most cancers harbor intra-tumoral hypoxia which promotes tumor progression and therapy resistance. Hypoxia-inducible factor 1α (HIF1α) mediates an adaptive response to hypoxia and contributes to multiple cancer hallmarks. We describe cancer therapeutic targeting of HIF1α by combination of CDK4/6 inhibitors (CDK4/6i) and heat-shock protein 90 inhibitors (HSP90i). CDK1 contributes to HSP90-mediated HIF1α stabilization whereas CDK1-knockdown enhances HIF1α reduction by HSP90i. Dual CDK1- and HSP90-inhibition increases apoptosis and synergistically inhibits cancer cell viability. To translate our findings, we use FDA-approved CDK4/6i in combination with HSP90i to reduce HIF1α expression and suppress viability of multiple cancer cell types, including Rb-deficient cancer cells. Overexpression of HIF1α^668E^ partially rescues the cell viability inhibition by combination CDK4/6i and HSP90i treatment under hypoxia. CDK4/6i and HSP90i suppresses tumor growth *in vivo*. Thus, combined targeting of CDK4/6 and HSP90, through a drug class effect, inhibits HIF1α and shows preclinical anti-cancer therapeutic efficacy, including with Rb-deficiency.

## Introduction

Accompanying the unrestrained proliferation of malignant cells, solid tumors are generally deprived of an adequate oxygen supply^1^. Regions located further than the oxygen diffusion limit (~100 μm)^2^ to blood vessels become hypoxic. Hypoxia is implicated in cancer, linked to abnormal vascularization, altered metabolism, resistance to chemo-/radio-therapy, as well as increased cancer cell stemness and metastasis^3–7^. In adaptation to hypoxia, hypoxia-inducible factor 1 (HIF1), as a transcription factor, stimulates a variety of target genes that are involved in altered metabolism, cell survival and tumor progression^8–10^. In particular, the α subunit of HIF1, HIF1α, becomes constitutively expressed, which leads to the constant activation of HIF1.

Overexpression of HIF1α is observed in a variety of cancers. In colorectal cancer (CRC), it is associated with poor prognosis and early progression^11,12^. HIF1α inhibits apoptosis^13,14^, facilitates cell migration^14^ and promotes angiogenesis through upregulation of the target *VEGF* gene^15^ in CRCs and other tumors. When oxygen is sufficient, in normal cells, HIF1α is hydroxylated by prolyl hydroxylase-domain proteins (PHDs) and is targeted by the von Hippel-Lindau (VHL) protein complex for ubiquitination and subsequent proteasomal degradation^16^, which is prevented by hypoxia^17^. However, elevated HIF1α expression is not exclusive to hypoxic conditions. In renal cell carcinomas, VHL is frequently mutated and deficient^18^. In RCC4 renal cancer cells, for instance, HIF1α is constantly expressed at increased levels due to protein stabilization. EGF/EGFR signaling transcriptionally activates HIF-1α independently of hypoxia^19^. Moreover, HIF1α was shown to be detectable at other regions in the tumor other than the hypoxic necrotic margin^20^. HIF1α can accumulate in T_H_17 cells under normoxia and regulates T_H_17 differentiation, suggesting a role of HIF1α in the immune system in both normoxia and hypoxia^21^.

We previously carried out a chemical library screen for hypoxia sensitizers and uncovered cyclin-dependent kinase inhibition as a potential therapeutic strategy for hypoxic tumors^22^. We further showed that cyclin-dependent kinase 1 (CDK1) stabilizes HIF1α through phosphorylation of the Ser668 residue of HIF1α protein^23^. Such stabilization occurs not only in hypoxia, but also at the G2/M cell cycle phase under normoxic conditions. Moreover, CDK4 is also important for HIF1α stabilization, as we uncovered in our study through knockdown of CDK proteins^23^. Assessment of RCC4 cells demonstrated that the CDK1- or CDK4-inhibitor-mediated HIF1α destabilization is independent of a functional VHL protein.

Another VHL-independent HIF1α stabilizer is the heat shock protein 90 (HSP90)^24^. HSP90 is a HIF1α-associated protein^25^. Overexpression of HSP90 has been correlated with adverse prognosis and recognized as a therapeutic target in cancer (e.g. esophageal squamous cell carcinoma, melanoma, leukemia) ^26–28^. Both CDK and HSP90 inhibitors have been widely studied^29,30^. Ro-3306 is a CDK1-selective inhibitor^31^. The CDK4/6 inhibitor, palbociclib, among others has been approved by the FDA in combination treatment for breast cancer^32,33^. HSP90 inhibitors have evolved from the classical small molecule geldanamycin to second generation compounds (e.g. ganetespib^34^). In colorectal cancer, ganetespib has been found to inhibit angiogenesis^35^ and sensitizes cells to radiation and chemotherapy^36^.

We initially investigated the hypothesis that CDK1 and HSP90 signaling overlaps in the regulation of HIF1α, and that combination treatment to target both CDK1 and HSP90 may lead to enhanced inhibitory effects towards HIF1α expression and function as well as improved anti-cancer efficacy. We extended our observations to CDK4/6 inhibitors given the fact there are several FDA-approved drugs allowing more rapid translation of our findings. We uncovered a synergy between CDK4/6 inhibitors and HSP90 inhibitors, as a class effect for each of the two drug classes, through convergence upon HIF1α leading to cell death. Importantly, the dual blockade of CDK4/6 and HSP90 is observed in Rb-deficient tumor cells suggesting a novel approach for cancer therapy. We investigated overexpression of HIF1α^668E^, a mimic of phosphorylated HIF1α, and showed that it partially rescues cell viability inhibition by combined CDK4/6i and HSP90i treatment under hypoxia. An additional aspect of this work involves a focus on HIF as a potential biomarker for CDK4/6-HSP90 dual inhibition therapy. Overall, our results suggest a novel therapy combination that is efficacious in preclinical models for targeting hypoxic tumor cells and this could be further advanced and tested in clinical trials.

## Results

### CDK1 contributes to HSP90-mediated HIF1α stabilization

We have previously reported that knockdown of CDK1 led to the reduction of HIF1α level in RCC4 VHL-deficient cells^23^. To reinforce the hypothesis that the regulatory effect on HIF1α by CDK1 inhibition is independent of VHL, we examined the level of HIF1α upon addition of CDK1 inhibitor, Ro-3306, in both RCC4 and RCC4^VHL+^ cells. As expected, HIF1α was constantly expressed in RCC4 cells under normoxia, owing to the loss-of-function mutation of *VHL* in this cell line. In accordance with previous results, CDK1 inhibition reduced HIF1α level in RCC4 in normoxia, which could be reversed by proteasome inhibition with MG132 (Fig. 1A). In RCC4^VHL+^ cells where VHL is stably reintroduced, HIF1α expression was dramatically decreased in normoxia compared to that in RCC4 cells. Inhibition of CDK1 decreased the level of HIF1α in RCC4^VHL+^ cells, which could be rescued with MG132 (Fig. 1A). Thus, CDK1 inhibition destabilized HIF1α in a VHL-independent manner.

**Figure 1.**
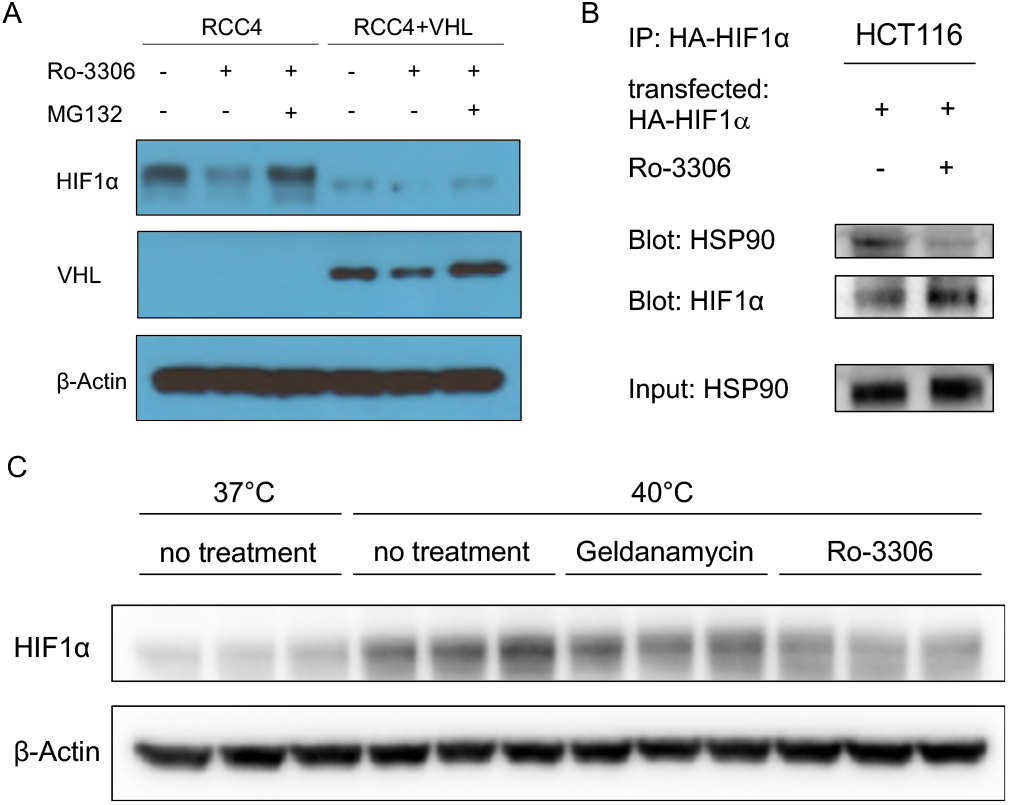
CDK1 contributes to HSP90-mediated HIF1α stabilization. (A) Inhibition of CDK1 decreases the level of HIF1α in RCC4 cells independently of VHL. Cells were treated with Ro-3306 (5 μM) or MG132 (1 μM) or both as indicated for 6 hours under normoxia. (B) CDK1 inhibition (for 6 hours under hypoxia; 0.5% O_2_) impairs the interaction between HIF1α and HSP90. HCT116 cells were treated with MG132 and cultured in hypoxia for 6 hours with or without Ro-3306. Cells were fixed and lysed for co-immunoprecipitation analysis. (C) CDK1 partially reversed heat shock-induced HIF1α expression. HCT116 cells were treated at 40°C with the indicated inhibitors for 6 hours.

Another previously known VHL-independent HIF1α stabilizer and associating partner is HSP90^ref24,25^. We asked whether there is a link between CDK1-mediated and HSP90-mediated stabilization of HIF1α. We found that inhibition of CDK1 impaired the interaction between HIF1α and HSP90 (Fig. 1B). Moreover, heat shock (40°C) induced HIF1α expression in normoxia, which could be partially reversed by treatment with HSP90 inhibitor, geldanamycin, or CDK1 inhibitor, Ro-3306 (Fig. 1C). These results suggest that CDK1 may contribute to the stabilization of HIF1α by HSP90.

### Dual targeting of CDK1 and HSP90 robustly reduces the expression level of HIF1α

On basis of the findings above, we tested whether targeting CDK1 could enhance the inhibitory effect on HIF1α expression by HSP90 inhibitors. Consistent with previous findings, the level of hypoxia-induced HIF1α was decreased by CDK1 knockdown or HSP90 inhibition with geldanamycin. Remarkably, when geldanamycin was added to CDK1-knockdown cells, the reduction of HIF1α was further enhanced (Fig. 2A). Such enhanced HIF1α inhibition was also observed with combination treatment using the two inhibitors, Ro-3306 and geldanamycin (Fig. 2B).

**Figure 2.**
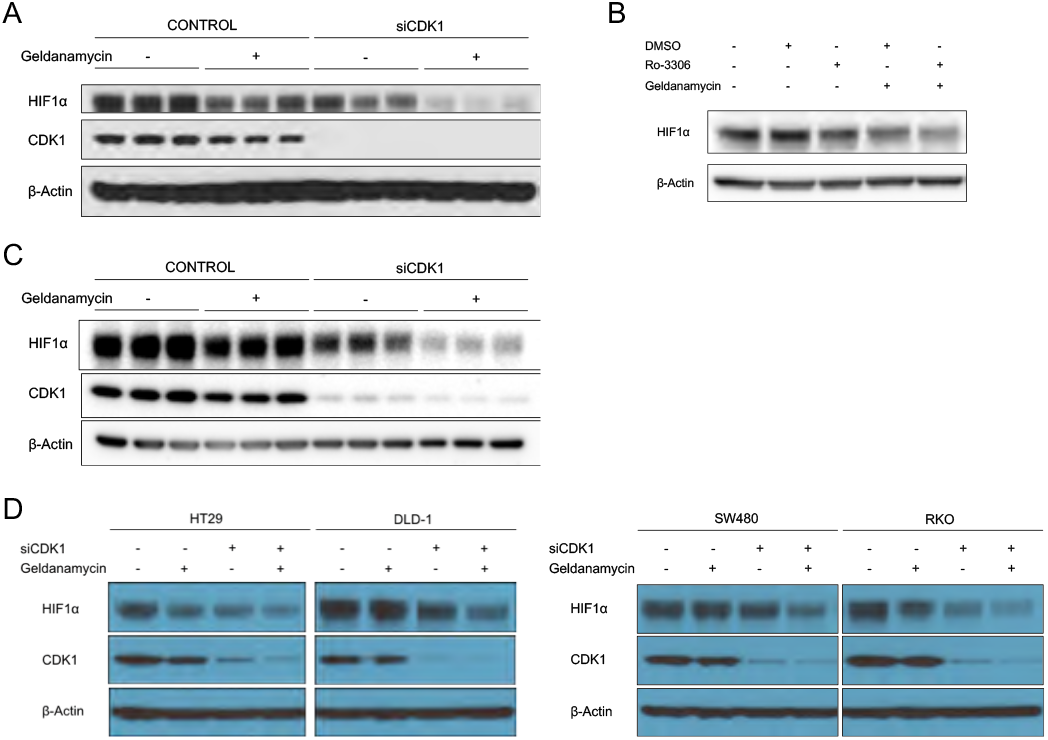
Dual inhibition of CDK1 and HSP90 robustly reduces the level of HIF1α. (Α) HCT116, (C) HCT116 p53−/− cells or (D) other colorectal cell lines were treated with control or CDK1 siRNA for 48 hours, followed by treatment with DMSO or geldanamycin under hypoxia (0.5% O_2_) for 6 hours. (B) Cells were treated with Ro-3306, geldanamycin, or the combination of both for 6 hours under hypoxia.

It is known that p53 is mutated in approximately 40%-50% of sporadic colorectal cancers^37^, we tested whether the absence of p53 affects the combinational effect. In HCT116 p53^−/−^ cells, combination treatment robustly diminished the level of HIF1α similarly as in wild-type cells (Fig. 2C), indicating that the HIF1α-regulatory effect is p53-independent. Consistently with these observations, the enhanced inhibition of HIF1α by combination treatment was observed in other colorectal cancer cells with different p53 status (Fig. 2D) (i.e. HT29: p53^G273A^; DLD1: p53^C241T^, SW480: p53^G273A&C309T^; RKO: p53^wild-type^).

### Dual inhibition of CDK1 and HSP90 synergistically suppresses cancer cell viability

The universal effect of HIF1α inhibition by combination of CDK1 knockdown and HSP90 inhibition among various colorectal cancer cell lines prompted us to investigate the therapeutic potential of such combination strategy. We performed a CellTiter-Glo assay to assess the combinatorial effect on cell viability by CDK1 and HSP90 inhibitors. We found that Ro-3306 and geldanamycin synergistically inhibited HCT116 cell viability in both normoxia and hypoxia (Fig. 3A, B).

**Figure 3.**
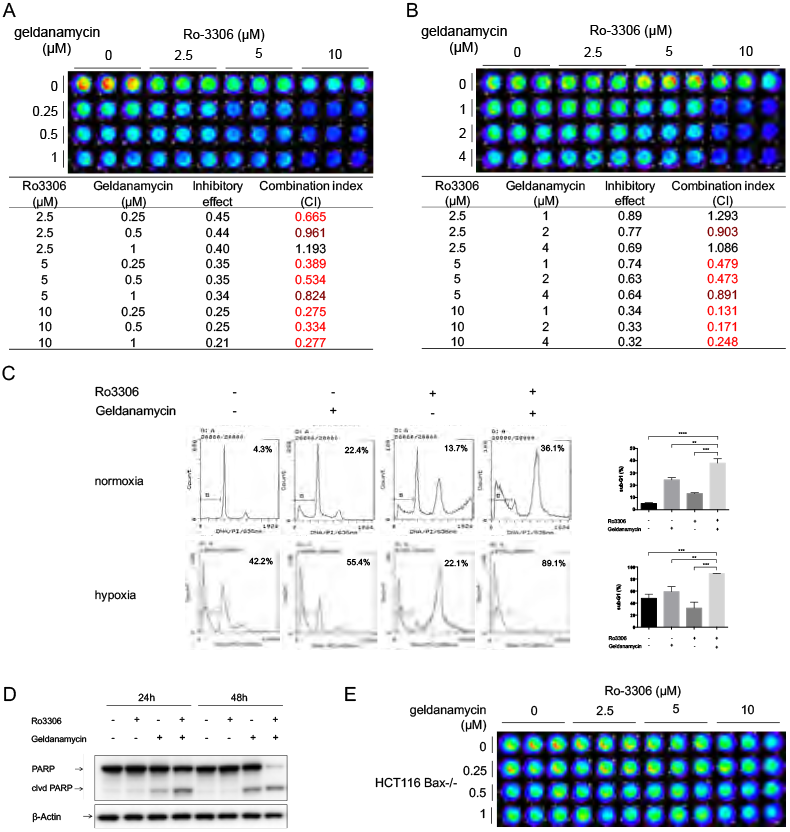
Ro-3306 and geldanamycin synergistically inhibit HCT116 cell viability through induction of apoptosis. (A) In normoxia or (B) hypoxia (0.5% O_2_), cells were treated with Ro-3306 and geldanamycin at the indicated concentrations for (A) 48 or (B) 72 hours. (C) Sub-G1 analysis by propidium iodide staining and flow cytometry of cells treated with Ro-3306 (10 μM) and geldanamycin (1 μM) under normoxia for 48 hours or under hypoxia for 72h. (D) Western blot of PARP cleavage in cells treated with Ro-3306 or geldanamycin or both. (Ε) CellTiter-Glo analysis of cell viability in HCT116 Bax−/− cells treated at indicated concentrations under normoxia for 48 hours. ** p<0.01; *** p<0.005; **** p<0.001.

Subsequently we asked whether apoptosis was induced by the combination treatment. We performed sub-G1 analysis by flow cytometry to estimate fractional DNA content^38^. Combination of Ro-3306 and geldanamycin significantly increased the sub-G1 population in HCT116 cells as compared to control and single treatments in either normoxia or hypoxia (Fig. 3C), indicating the increased occurrence of apoptosis. As expected, PARP cleavage was also observed (Fig. 3D) at an earlier time point as a marker of initiated apoptosis^39^. In addition, the robust synergy on cell viability inhibition was abrogated in HCT116 Bax^−/−^ cells (Fig. 3E), indicating that Bax may play an important role in mediating cell death induced by the CDK1i/HSP90i combination treatment.

### Dual inhibition of CDK1 and HSP90 represses the ability of colony formation and cell migration

Not every single cancer cell is capable of proliferating into a colony^40^. To determine whether the combination treatment as well induces cell reproductive death in an *in vitro* model, we performed clonogenic assays to test the post-treatment change in cell capability to generate colonies. Treatment with both Ro-3306 and geldanamycin, at relatively low doses (2.5 μM, 0.02 μM, respectively), markedly inhibited colony formation of HCT116 cells in both normoxia (Fig 4A) and hypoxia (Fig. 4B). The colonies that formed upon combination treatment were fewer in number and smaller in size as compared to control and single treatments. Thus, the dual inhibition of CDK1 and HSP90 inhibits colony formation by HCT116 colon cancer cells.

**Figure 4.**
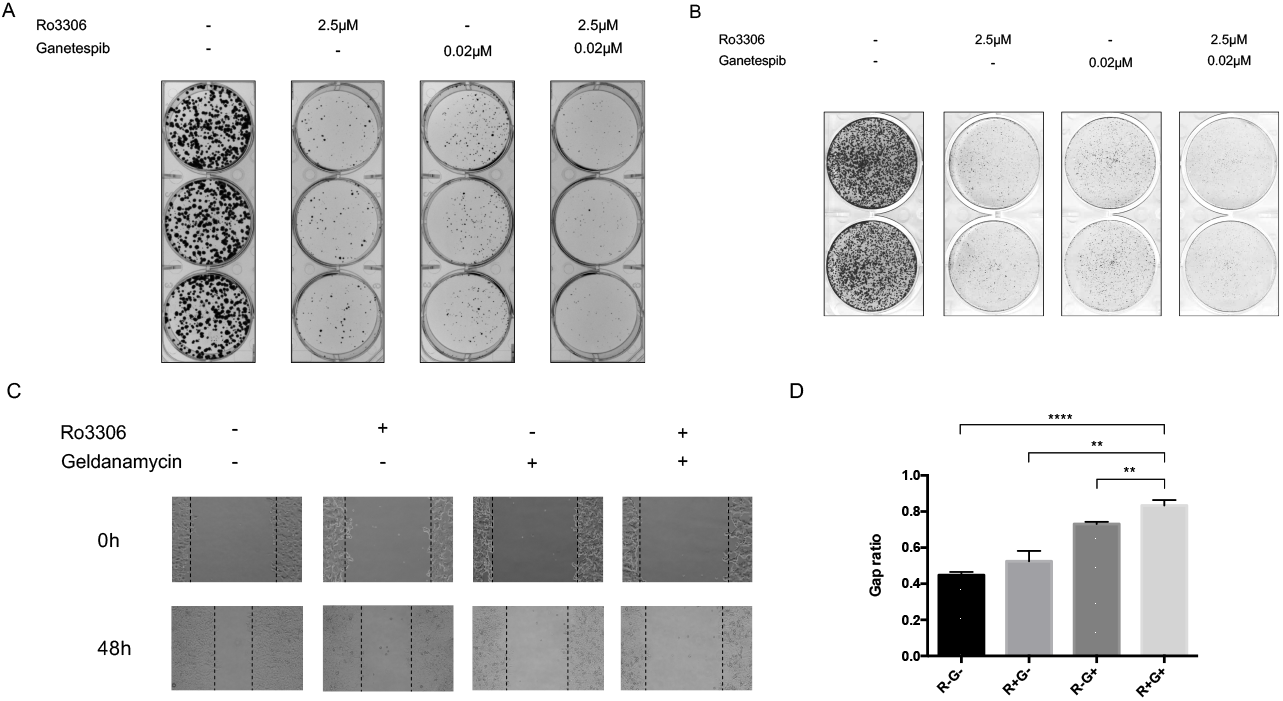
Combination CDK1 and HSP90 inhibitor treatment inhibits colony formation and migration in HCT116 cells. (A) In normoxia or (B) In hypoxia (0.5% O_2_), HCT116 cells were treated with the indicated combination treatments for 72 hours. Drug-containing media was replaced with regular culture media, and cells were allowed to grow and form colonies for 1 week. (C) Scratch assay and (D) quantification for HCT116 cells under normoxia for 48 hours. Gap ratio refers to the ratio of gap width at 48 hours versus at 0 hours. Cells were treated with Z-VAD caspase inhibitor to prevent cell death. (R: Ro-3306; G: geldanamycin.) n=3. ** p<0.01; *** p<0.005; **** p<0.001.

The overexpression of HIF1α in cancer is implicated not only in promoting cell survival but also in cell migration^41^. We performed an *in vitro* scratch assay^42^ to test the effect of combination treatment on HCT116 motility. An artificial gap was created on a nearly confluent monolayer of cells. The cell monolayer bearing wounds was treated with single or combination of the two inhibitors, together with Z-VAD-FMK, a pan-caspase inhibitor to prevent treatment-induced cell death. Gap ratio was calculated using gap width at 48 hours normalized to that at 0 hour. The ratio was significantly higher in the combination treatment group as compared to the control and single treatment groups (Fig. 4C, D), suggesting that the combination of Ro-3306 and geldanamycin inhibits HCT116 cell migration.

### Dual inhibition of CDK4/6 and HSP90 shows anti-cancer effects

We have previously shown that knockdown of CDK4 was able to reduce the level of HIF1α^23^. Considering the clinical use of the FDA-approved CDK4 inhibitors, we sought to examine the anti-cancer effects by CDK4 inhibition in combination with HSP90 inhibitors. Two different HSP90 inhibitors, ganetespib and onalespib, were tested first in the study. As expected, either ganetespib or onalespib alone reduced the expression level of HIF1α (Fig. 5A, B). The addition of CDK4 inhibitor, palbociclib, was able to further enhance the HIF1α decrease induced by HSP90 inhibition (Fig. 5A, B). Knockdown of CDK4 with siRNA exhibited a similar effect (Supplementary Fig. 1A). Combination treatment with palbociclib and either of the HSP90 inhibitors showed synergistic inhibition on cell viability in HCT116 cells in both normoxia and hypoxia (Fig. 5C, D). Such synergy was also observed in other colorectal cancer cells (e.g. SW480, Supplementary Fig. 1B, C). Dual inhibition of CDK4 and HSP90 significantly increased the sub-G1 population in HCT116 cells regardless of oxygen concentration (Fig. 5E). Combination treatment with palbociclib and ganetespib significantly inhibited HT29 cell migration in CoCl_2_-treated cells where hypoxia is mimicked (Fig. 5F). These results indicate that targeting CDK4/6 in combination with HSP90 inhibition has a similar anti-cancer effect as dual inhibition of CDK1 and HSP90.

**Figure 5.**
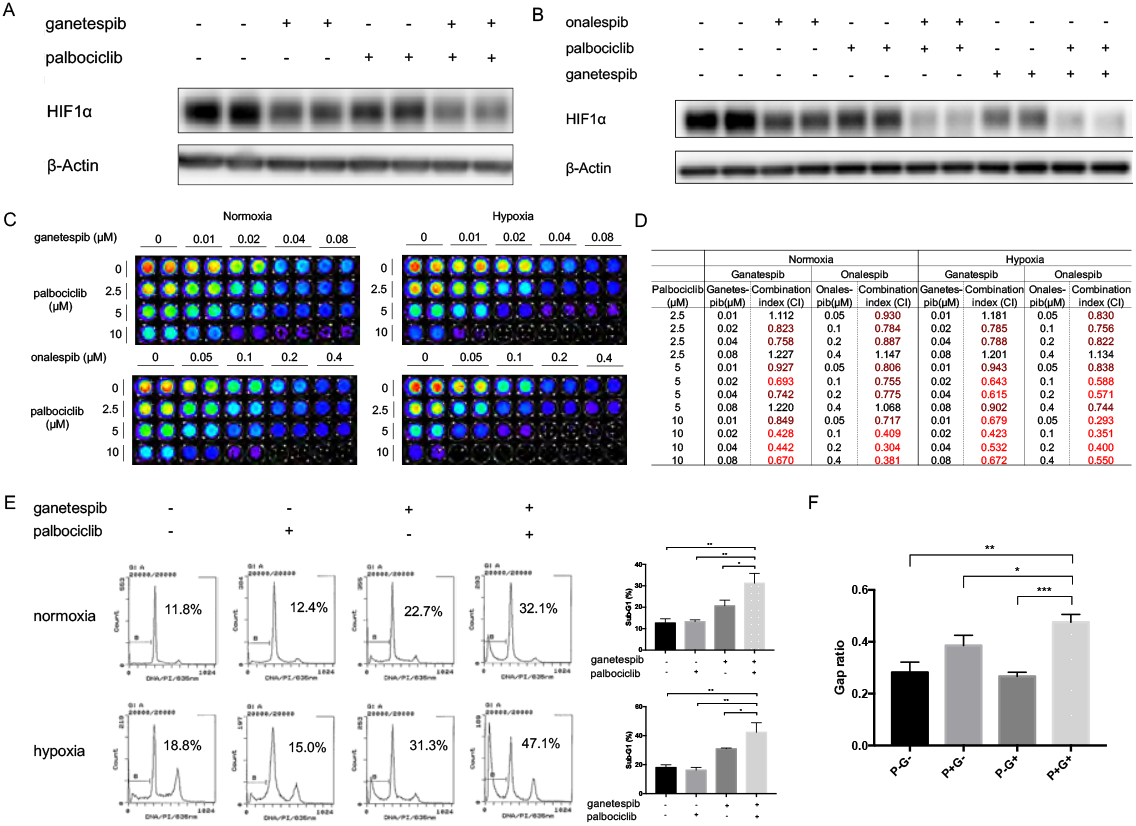
Dual inhibition of CDK4/6 and HSP90 reduces HIF1α in colorectal cancer cells and synergistically inhibits cell viability in HCT116. (A) HCT116 cells were treated with the indicated inhibitors (ganetespib at 0.05 μM and palbociclib at 10 μM) for 6 hours under hypoxia (0.5% O_2_). (B) SW480 cells were treated with the indicated inhibitors (ganetespib at 0.05 μM, onalespib at 0.05 μM, and palbociclib at 10 μM) for 6 hours under hypoxia (0.5% O_2_). (C, D) CDK4 inhibitor palbociclib and HSP90 inhibitor ganetespib or onalespib synergistically inhibit the viability of HCT116 cells at 72 hours in normoxia and hypoxia (0.5% O_2_). (E) Sub-G1 analysis by propidium iodide staining and flow cytometry for HCT116 cells treated with the indicated drug combinations (ganetespib at 0.04 μM; palbocilib at 10 μM) for 48 hours. (F) Scratch assay in HT29 cells under CoCl_2_ treatment (50μΜ) to mimic hypoxia for 72 hours. (P: palbociclib; G: ganetespib.) * p< 0.05; ** p<0.01; *** p<0.005.

CDK4/6 inhibitors have been intensively studied in combination therapies. After palbociclib, two CDK4/6 inhibitors, ribociclib and abemaciclib, were approved as anti-cancer drugs. Meanwhile there have been many efforts in developing HSP90 inhibitors intended for cancer treatment with tolerable toxicity. To further test the translational potential of the dual inhibition strategy, we included the CDK4/6 inhibitor abemaciclib and two other HSP90 inhibitors that were being examined in clinical trials, XL-888 and TAS-116, in this study. Consistent with the results above, XL-888, in combination with palbociclib, exhibited similar inhibitory effects on HIF1α expression as well as cell viability (Supplementary Fig. 2). The combination of TAS-116 and palbociclib or abemaciclib markedly reduced the level of HIF1α (Supplementary Fig. 3A, D) and synergistically suppressed cell viability in SW480 colon cancer cells both in normoxia (Supplementary Fig. 3B, E) and hypoxia (Supplementary Fig. 3C, F).

These results not only established the preclinical foundation for potentially testing these drugs in clinical trials, but further confirmed a class effect of CDK4/6 and HSP90 dual inhibition in colorectal cancer treatment.

### Anti-tumor efficacy *in vivo* by combination treatment with palbociclib and ganetespib

To determine the anti-tumor efficacy of CDK4/6 and HSP90 dual inhibition *in vivo*, we used HT29 cancer cells in a xenograft mouse model. HT29 is relatively resistant to ganetespib compared to other colorectal cancer cell lines (Supplementary fig. 4). We tested whether the addition of palbociclib could improve the tumor-suppressive performance of ganetespib. The weight of drug combination-treated tumors was significantly lower than that of control and single treatment groups (Fig. 6A, B). Relative tumor volume was also low in the combination treatment group (Fig. 6C). There was no evident toxicity or weight loss observed upon the combination treatment compared to the control group (Fig. 6D), indicating the safety of simultaneous administration with palbociclib and ganetespib.

**Figure 6.**
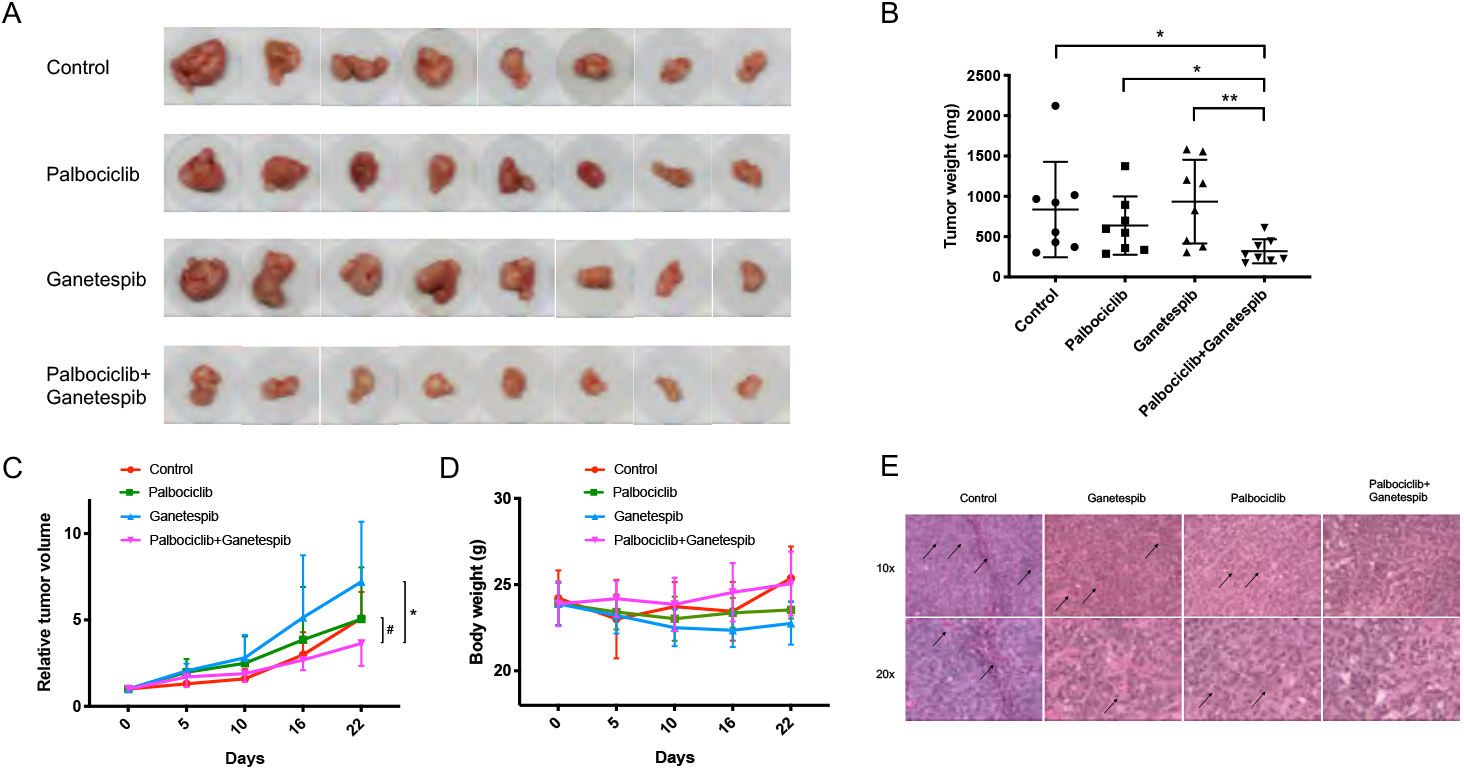
Combination treatment with palbociclib and ganetespib inhibits tumor growth *in vivo*. (A) Tumors excised from HT29 xenografts in nude mice. (B) Tumor weight quantification of excised tumors. (C) Relative tumor volume measured over time. Tumor volumes were normalized to those at the beginning of treatment. (D) Body weight of mice in different treatment groups. (E) Combination treatment inhibits microvessel formation in tumors *in vivo*. # control *vs*. combination: p<0.05; * ganetespib *vs*. combination: p<0.05.

Interestingly, the combination treatment reduced the presence of microvessels in tumors (Fig. 6E), which is consistent with suppression of the role of HIF1α in angiogenesis. In addition, the combination treatment increased caspase 3 cleavage and inhibited VEGF expression in the xenografts (Supplementary fig. 5). The *in vivo* results suggest a therapeutic potential of the CDK4/6 and HSP90 dual inhibition strategy in cancer treatment.

### Combination treatment of CDK4/6 and HSP90 inhibitors synergistically inhibit cell viability in multiple cancer types

Although the dual inhibition was mainly evaluated in colorectal cancers in this study, the strategy is not necessarily limited to one cancer type. CDK4/6 inhibition was initially investigated and approved for treatment in breast cancers. Hypoxia is a prominent characteristic of the tumor microenvironment in pancreatic cancer and glioblastoma, both of which lack efficacious treatments. Thus, we tested the effect of CDK4/6 and HSP90 dual targeting on HIF1α expression in various cancer cell lines. In our later studies, we have focused on using TAS-116 as the HSP90 inhibitor as it is currently being tested in early phase clinical trials for cancer. Enhanced HIF1α inhibition was shown upon the combination treatment of palbociclib and TAS-116 in ASPC1 and HPAFII pancreatic cancer cell lines (Supplementary fig. 6A, B) as well as SKBR3 and MDA-MB-361 breast cancer cells (Supplementary fig. 6C, D). Palbociclib and TAS-116 synergistically inhibited SKBR3 cell viability in both normoxia and hypoxia (Supplementary fig. 6E, F). Moreover, ganetespib and palbociclib diminished HIF1α expression in T98G glioblastoma cells (Supplementary fig. 7A). We have also found that knockdown of CDK4 in combination with HSP90 inhibition inhibited the level of HIF1α in PC3 prostate cancer cell line (Supplementary fig. 7B). These findings suggest that it may be worthwhile to pursue the translational potential of such combination treatment in more cancer types. We are currently pursuing a novel phase 1b clinical trial combining palbociclib with TAS116 in patients with breast cancer and other solid tumors.

### Rb-deficiency does not block the combinatorial inhibition of HIF1α and reduced cancer cell viability due to targeting of CDK4/6 and HSP90

Rb is a key downstream factor of CDK4/6 activity in cell cycle regulation. Loss of Rb protein is believed to convey resistance to CDK4/6 inhibitors. Here we tested whether the inhibitory effect by the combination treatment was diminished by Rb-deficiency. Saos2 is an osteosarcoma cell line which is naturally Rb-deficient. The combination treatment with abemaciclib and TAS116 synergistically inhibited cell viability at different doses in Saos2 cells in normoxia and hypoxia (Fig. 7A, B). We also knocked down Rb in Rb-proficient (wild-type) cell lines. Knockdown of Rb in SW480 cells and MCF7 cells did not affect the inhibitory effect on HIF1α expression upon combination treatment (Fig. 7C, D). The combination treatment also showed synergistic inhibition of cell viability in Rb-knockdown SW480 cells (Fig. 7E, F).

**Figure 7.**
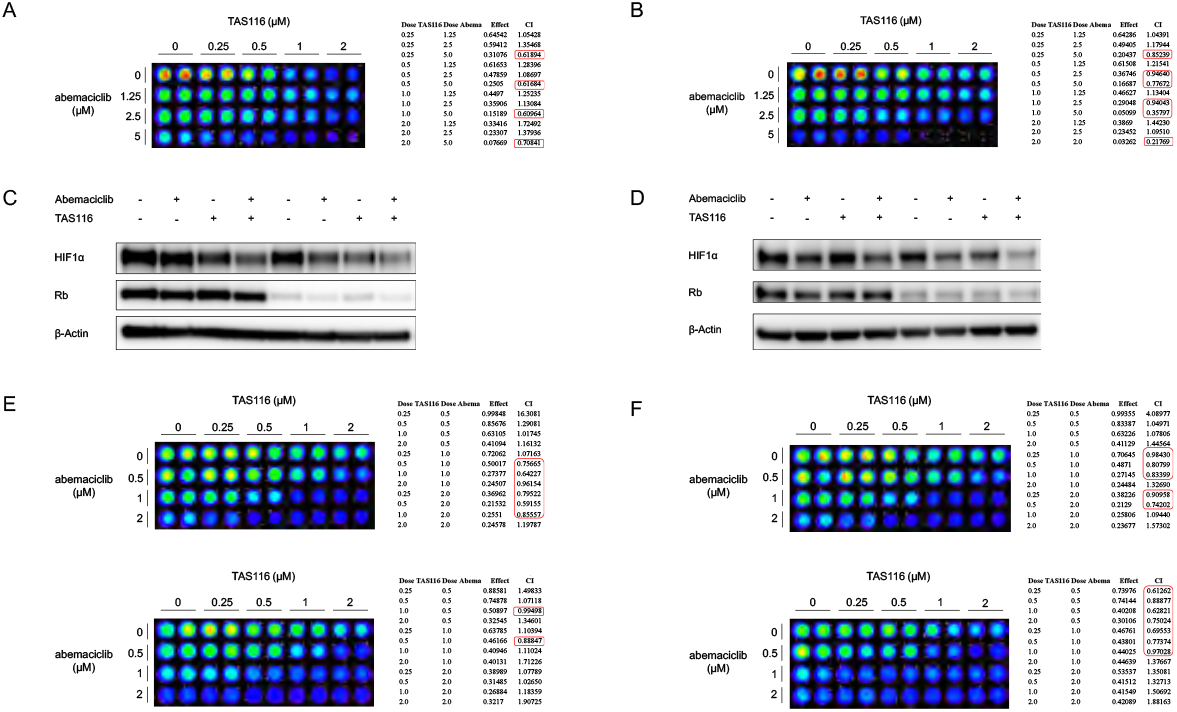
Rb-deficiency does not affect the combinatorial inhibition of HIF1α expression and cell viability. (A, B) Combination treatment with abemaciclib and TAS116 synergistically inhibits cell viability in Saos2 osteosarcoma cells at 72 hours under (A) normoxia and (B) hypoxia (0.5% O_2_). (C) SW480 cells were incubated with Rb-targeting siRNA for 48 hours and subsequently treated with 1 μM abemaciclib and/or 0.5 μM TAS116 for 6 hours in hypoxia (0.5% O_2_). (D) Knockdown of Rb does not affect HIF1α inhibition by combination drug treatment with TAS116 and abemaciclib in MCF7 breast cancer cells. (E, F) SW480 cells were treated with (E) mock or (F) Rb-targeting siRNA for 48 hours and subsequently treated with indicated combinations under normoxia (upper) or 0.5% O_2_ hypoxia (lower).

### HIF1a targets VEGFA and SLC2A1 correlate with poor disease-free prognosis in colorectal cancer

HIF1α is involved in multiple key signaling pathways in cancer progression. We analyzed the TCGA database on colon and rectal cancer using UCSC Xena online exploration tool. The overexpression of HIF1α target genes *VEGFA* and *SLC2A1* correlated with poor disease-free prognosis in colorectal cancer (Fig. 8). In clinic, pancreatic cancer is often highly hypoxic. The overexpression of HIF1α target genes *SLC2A1* and *PPIA* are associated with poor prognosis in pancreatic adenocarcinoma (Supplementary fig. 12A). Thus, targeting HIF1α may serve as a promising modality in cancer treatment as the poor prognostic factors would be inhibited by the proposed dual CDK4/6 and HSP90 inhibition strategy. Both HIF1α and its targets could serve as useful biomarkers in future clinical trials of dual CDK4/6 and HSP90 inhibitor therapy.

**Figure 8.**
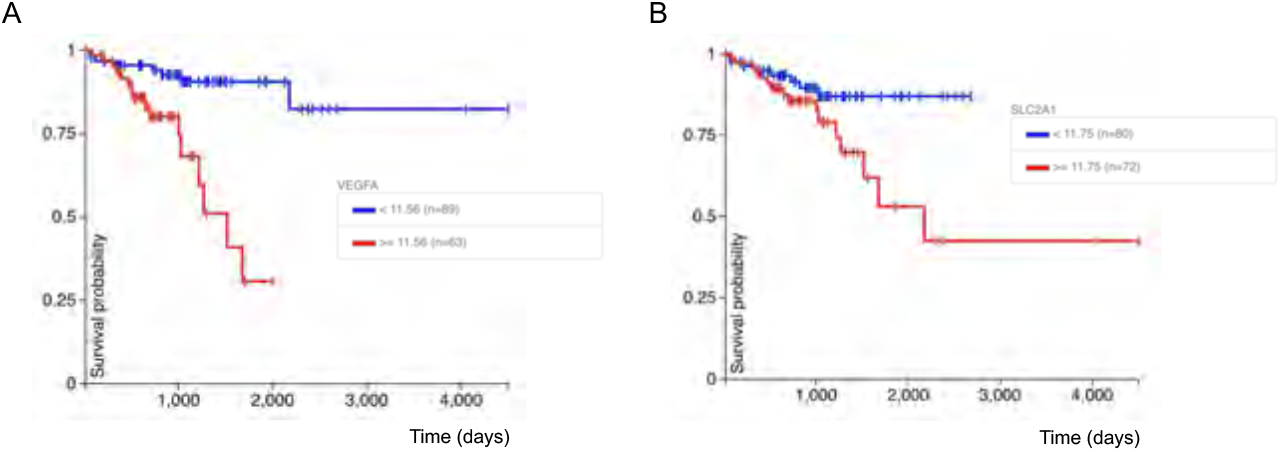
Correlation between the overexpression of HIF1α target genes *VEGFA* and *SLC2A1* and poor disease-free interval. The analysis was performed with the UCSC Xena tool on TCGA colon and rectal cancer samples. (A) Correlation between *VEGFA* expression and disease-free interval. P value=0.0001. (B) Correlation between *SLC2A1* expression and disease-free interval. P value=0.034.

## Discussion

We demonstrate a novel convergence of CDK4/6 and HSP90 dual inhibition on HIF1α inhibition that is VHL-, p53-, or hypoxia-independent and which can be translated as a cancer therapy, including for tumors with Rb-deficiency. In this regard, the data in this manuscript provides the preclinical rationale for a planned clinical trial combining CDK4/6 inhibitor palbociclib with HSP90 inhibitor TAS-116. The trial is planned for patients with breast cancer who have progressed on CDK4/6 inhibitor therapy and for patients with other solid tumors that are Rb-deficient. The patent by Zhao S. and El-Deiry W.S., “Dual Inhibition of CDK and HSP90 Destabilizes HIF1α and Synergistically Induces Cancer Cell Death”, US Patent 10,729,692 issued on August 4, 2020.

Hypoxia and HIF1α contribute to the malignant cancer progression phenotype across diverse cancer types. HIF1α is hyperactivated and participates in promoting breast cancer progression^43,44^. Anabolic metabolism induced by HIF1α leads to gemcitabine resistance in pancreatic cancer^45^. Also, hypoxia/HIF1α exerts a tumor-promoting role by immunosuppression. HIF-1α/VEGF-A signaling is indispensable for the tumor infiltration and cytotoxicity of effector CD8^+^ T cells in breast cancer^46^. Depletion of HIF1α in natural killer (NK) cells disturbs angiogenesis and inhibits tumor growth in the MC38 (colon cancer) isograft mouse model^47^. The immune checkpoint protein PD-L1 has been identified as a direct target of HIF-1α^48^. Meanwhile the pro-cancer effect by hypoxia is not limited to solid tumors. Indeed, the local oxygen tension appears quite low in bone marrow *in vivo*^49^. Hypoxia/HIF1α signaling maintains leukemia stem cells^50^ and facilitates invasion and chemo-resistance^51^ in T-ALL. It may be useful in cancer therapy to pursue effective strategies of targeting hypoxia and HIF1α signaling.

On the basis of our previous findings showing CDK1-mediated stabilization of HIF1α and also with the established role of HSP90 in HIF1α expression, we hypothesized a model where CDK1 contributes to HSP90-mediated stabilization of HIF1α. In our present studies, dual targeting of CDK1 or CDK4/6 and HSP90 robustly reduced the level of HIF1α and synergistically inhibited cell viability in colorectal cancer lines. To assess the anti-tumor effect, the combination of palbociclib and ganetespib was tested on HT29 xenografts. Palbociclib has been used in a colon carcinoma xenograft model at the dose up to 150 mg/kg p.o. once per day to achieve tumor burden suppression^52^. Ganetespib has been used in a HCT116 xenograft colon cancer model at 150 mg/kg i.v. once per week to inhibit tumor growth^36^. In the present study, we administrated into the mice *considerably lower doses* of both compounds (palbociclib at 50 mg/kg; ganetespib at 25 mg/kg). We expect for therapeutic purposes, there would be less toxicity associated with HSP90 inhibition by reduced dosing in this strategy. As the result showed, body weights were not affected by the combination therapy compared to control. However, this does not necessarily preclude the possibility of increasing the doses of each drug in case they are well-tolerated. In the relative tumor volume measurement (Fig. 6C), although an inhibitory trend was shown by combination treatment, no significant difference was indicated by statistical analysis between the palbociclib alone and the combination groups. This may be due to the accuracy of measurements, variation among individual subjects and limited numbers of animals per group. Notably, HT29 is a relatively resistant cell line to ganetespib. The combination with palbociclib sensitized the xenografts for ganetespib treatment. Combination of palbociclib and ganetespib did not trigger synergistic toxicity to WI38 normal cells in normoxia *in vitro* (Supplementary fig. 8). Recently, efforts have been made to develop new generation of HSP90 inhibitors, which may contribute alternative choices other than ganetespib itself. Thus, we are planning to use HSP90 inhibitor TAS-116 in combination with palbociclib in a planned clinical study based on the rationale provided in this manuscript.

Due to the involvement of HIF1α in multiple aspects in cancer biology, whether the combination treatment affects other HIF1α-mediated cancer phenotypes remains to be tested. For instance, HIF1α plays an essential role in stem cell-induced target cell invasion^53^. Hypoxia/HIF1α can regulate cancer stem cell-like features^54,55^. It is not clear whether the CDKi (CDK inhibition) plus HSP90i (HSP90 inhibition) treatment modulates cancer stemness. Also, the effect of combination treatment on metastasis remains to be unraveled, considering the function of HIF1α as a driving force for metastasis/invasiveness^14,56–58^. In addition, since hypoxia/HIF1α is implicated in many immunosuppressive mechanisms^59–62^, it will be of interest to determine whether the combination CDKi/HSP90i treatment modulates the immune response for anti-tumor activities. CDK inhibition has recently been shown to stimulate tumor immune response^63–65^. Furthermore, it remains undefined whether any predictive biomarker(s) could be used to indicate sensitivity to the combination CDKi/HSP90i treatment. In this regard, HIF expression and HIF targets are prime candidate biomarkers. The enhanced inhibition of HIF1α by combined targeting of CDK1 or CDK4/6 and HSP90 was observed in multiple tumor cell lines. It would be useful to explore the anti-cancer effect of combination CDKi/HSP90i treatment in additional cancer types, and based on our results, we plan to include Rb-deficient solid tumors in the phase 1b study.

We performed a preliminary test on HIF2α expression. The combination treatment slightly reduced the level of HIF2α in HCT116 cells at 6 hours (Supplementary fig. 9). It may be interesting to investigate the effect on HIF2α according to its role in different cancer types (*e.g.* clear-cell renal cell carcinoma).

To test the involvement of HIF1α in the combination treatment, we transiently transfected HCT116 cells with plasmids containing HA alone or HA-HIF1α^668E^, a HIF1α mutant that remains stable upon CDK inhibition^23^. Overexpression of HIF1α^668E^ partially rescued the cell viability inhibition by combination treatment under hypoxia (Supplementary fig. 10), indicating that HIF1α may play a role in the combination effect. Since E2F signaling serves as an indicator of CDK4/6 activity, we performed a Pearson correlation test on some of the HIF1α and E2F target genes using the GEPIA tool based on TCGA colon adenocarcinoma data and pancreatic adenocarcinoma data, and found correlations between the expression of several HIF1α and E2F targets (Supplementary fig. 11 & Supplementary fig. 12B), which is consistent with the concept that CDK4/6 activity is linked to HIF1α signaling in patient tumors. As both Rb and HIF1α are molecular substrates for CDK4/6, we would suggest that HIF1α is a relevant and important target for CDK4/6 inhibitor therapy. In that context, HIF1α and its transcriptional targets may serve as useful biomarkers for drug efficacy, and the blockade of HIF1α may contribute to the anti-tumor effects of CDK4/6 inhibitors.

In summary, we provide a rationale for targeting HIF1α through a novel combination of CDK and HSP90 inhibitors as a potential therapeutic strategy. Our findings suggest new applications of previously approved CDK4/6 inhibitory drugs and novel HSP90 inhibitory agents in combination therapies in multiple cancer types including Rb-deficient tumors.

## Materials and Methods

### Cell culture

HCT116, SW480, HT29, DLD1 and RKO cells were obtained from American Type Culture Collection. HCT116, HT29 and SKBR3 cells were maintained in McCoy’s 5A medium (Hyclone) with 10% fetal bovine serum (FBS, Hyclone) and 1% penicillin/streptomycin (P/S). SW480, DLD1, RCC4, ASPC1, HPAFII, and T98G cells were maintained in Dulbecco’s modified Eagle medium (Hyclone) with 10% FBS and 1% P/S. RKO cells and PC3 cells were maintained in RPMI 1640 medium (Hyclone) with 10% FBS and 1% P/S. MDA-MB-361 cells were maintained in DMEM-F12 with 10% FBS, 1% P/S and 1% glutamine. Saos2 cells were maintained in McCoy’s 5A medium with 15% FBS and 1% P/S. WI-38 cells were maintained in Eagle’s Minimum Essential Medium with 10% FBS and 1% P/S. Cells were regularly tested for mycoplasma and authenticated. All cell lines were maintained at 37°C in 5% CO_2_. As for hypoxia treatment, cells were kept in a hypoxia chamber (In vivo2, Ruskinn) which maintains 0.5% O_2_.

### Antibodies and reagents

HIF1α and Ran antibodies were purchased from BD Biosciences. CDK1 and CDK4 antibodies were purchased from Santa Cruz Biotechnology. HA, Rb, HSP90, PARP and cleaved PARP antibodies were purchased from Cell Signaling Technology. Actin antibody was purchased from Sigma. HIF2α antibody was purchased from Novus Biologicals. MG-132 was purchased from Sigma. Ro-3306 was purchased from Santa Cruz Biotechnology. PD-0332991 (palbociclib) was purchased from Medkoo Biosciences. Geldanamycin was purchased from Invivogen. Ganetespib was purchased from ApexBio or Medkoo Biosciences. Onalespib was purchased from Cayman Chemical Company. XL888 was purchased from Medkoo Biosciences. TAS-116 was purchased from Active Biochem.

### Western blot

Treated cells were lysed in RIPA buffer (Sigma). Protein concentrations were determined using a BCA Protein Assay Kit (Life Technologies). Equal amounts of total protein were boiled with NuPAGE™ LDS sample buffer (Thermo Fisher Scientific) and reducing agent (Invitrogen) or 2-Mercaptoethanol. Samples were analyzed with SDS-PAGE. Proteins were transferred to an Immobilon-P PVDF membrane (EMD Millipore). Primary and secondary antibodies were added in order. Signals were detected after addition of the ECL western blotting substrate (Thermo Fisher Scientific).

### Cell transfection

Transient transfection of DNA was performed using Opti-MEM (Thermo Fisher Scientific) and Lipofectamine 2000 (Life Technologies). pcDNA3-HA-HIF1α plasmid was a gift from William Kaelin (Addgene plasmid #18949) ^66^. Knockdown experiments were performed with Opti-MEM and Lipofectamine RNAiMAX (Life Technologies), according to the manufacturer’s protocol. Control, CDK1 and CDK4 siRNAs were purchased from Santa Cruz Biotechnology. Rb siRNA was purchased from Cell Signaling Technology.

### Immunoprecipitation

HCT116 cells were transiently transfected with pcDNA3-HA-HIF1α. After 24 hours, cells were treated in hypoxia for 6 hours with MG132 (1 μM). Cells were washed with PBS and fixed in 4% formaldehyde. Cell lysis was performed in RIPA buffer with gentle sonication. The protein concentration in the lysates was measured and equalized. Part of the lysate was analyzed by SDS-PAGE and western blot for input monitoring. The remaining majority of the lysate was incubated with anti-HA antibody overnight at 4°C, followed by precipitation with Protein A/G Ultra link Resin (Thermo Fisher Scientific) for 2-4 hours.

### Synergy analysis

Indicated cells were seeded in a 96-well black microplate (Greiner Bio-One) and treated with combinations of inhibitors at various concentrations for 48 or 72 hours in normoxia or hypoxia. CellTiter-Glo reagent (Promega) was added and mixed on an orbital shaker at room temperature. Luminescence was recorded as a readout to compare viable cell number difference. Combination index between two treatments was calculated using Compusyn software. Synergism was indicated by a combination index value of < 1.

### Colony formation assay

Cells were seeded at the concentration of 500 cells/well in a 6-well plate and allowed to attach overnight. After subsequent drug treatment for 72 hours, the culture media was substituted with fresh drug-free complete media. Cells were kept in culture for one to two weeks with medium replacement every three days. At the endpoint, cells were rinsed with PBS and fixed with 10% formalin for 15 min. 0.05% crystal violet was used to stain the colonies. Plates were rinsed carefully in the sink with tap water and let dry at room temperature.

### Sub-G1 analysis

HCT116 cells were treated with indicated reagents for 48 or 72 hours in normoxia or hypoxia. Culture media including floating cells were collected and combined with trypsinized (Gemini Bio-Products) attached cells. All harvested cells were washed in PBS with 1% FBS. Cells were fixed with cold 70% ethanol at 4 °C. Subsequently, cells were washed, incubated in phosphate citrate buffer, and stained with propidium iodide (Sigma). The percentage of cells with sub-G1 DNA content was analyzed by propidium iodide staining and flow cytometry.

### Wound healing assay

The indicated cell lines were plated in 12-well plates at 80~90% confluence. Scratch lines were made with a 200-μL pipette tip. After washing with PBS, cells were cultured in media containing reagents as indicated. Images were captured at both the beginning and end of the experiment. Gap width was measured in each image. Each treatment group contained three replicates.

### *In vivo* studies

Animal experiments were conducted in compliance with the Institutional Animal Care and Use Committee at Fox Chase Cancer Center and followed the Guide for the Care and Use of Laboratory Animals. Hairless combined immunodeficient (SCID) mice were monitored in the Laboratory Animal Facility at Fox Chase Cancer Center. HT29 cells were subcutaneously injected into both rear flanks of 4-week old mice at 1×10^6^ / 100 μL in Matrigel/PBS. Treatments were started when tumors reached 100-125 mm^3^ as measured by Vernier caliper. Tumor-bearing mice were treated with palbociclib or ganetespib or the combination of both. Palbociclib was administered orally via gavage at 50 mg/kg daily (dissolved in ddH_2_O). Ganetespib was administered intravenously via retro-orbital injection at 25 mg/kg weekly (dissolved in 10% DMSO, 18% Cremophor RH 40, 3.8% dextrose). Growth of tumors was monitored for three weeks. At the endpoint, mice were euthanized, and tumors were dissected. The fixation, embedding (with Paraffin), sectioning and hematoxylin and eosin (H&E) staining of tumor samples were performed by the Histopathology Facility at Fox Chase Cancer Center.

### Statistical analysis

Results are presented as the mean ± standard deviation (SD). Difference comparisons were performed with Prism software using the Student’s two-tailed *t* test. Statistically significant differences were determined by P value < 0.05.

### Correlation analysis

Gene expression correlation analysis was performed using the GEPIA web server (http://gepia.cancer-pku.cn/) on colon adenocarcinoma TCGA data. Pearson correlation coefficient was calculated. The Kaplan-Meier plot was generated using UCSC Xena based on TCGA colon and rectal cancer (https://xena.ucsc.edu/) or using the GEPIA tool based on TCGA pancreatic adenocarcinoma cancer.

## Author contributions

S. Zhao participated in the conception, design, data acquisition and writing of the manuscript. L.Z. participated in the design and performance of *in vivo* study. D.T.D. participated in flow cytometry analysis. A.L. participated in helpful discussions. S. Zhang participated in discussion, provided technical advice and assisted with editing the manuscript. W.S.E-D. supervised this work and participated in the conception, design, analysis, writing and revision of the manuscript. W.S.E-D. provided oversight of the research as well as resources.

## Acknowledgements

We thank the El-Deiry lab members for all the helpful discussions. This work was presented in part as it progressed each year at the annual American Association for Cancer Research (AACR) meetings in 2016, 2017, 2018, 2019, and 2020. W.S.E-D. is an American Cancer Society (ACS) Research Professor and was supported by the William Wikoff Smith endowed professorship in cancer research at Fox Chase Cancer Center and by the Mencoff Family University Professorship in Medical Science at Brown University.

## Supplementary Figure Legends

**Supplementary figure 1.**
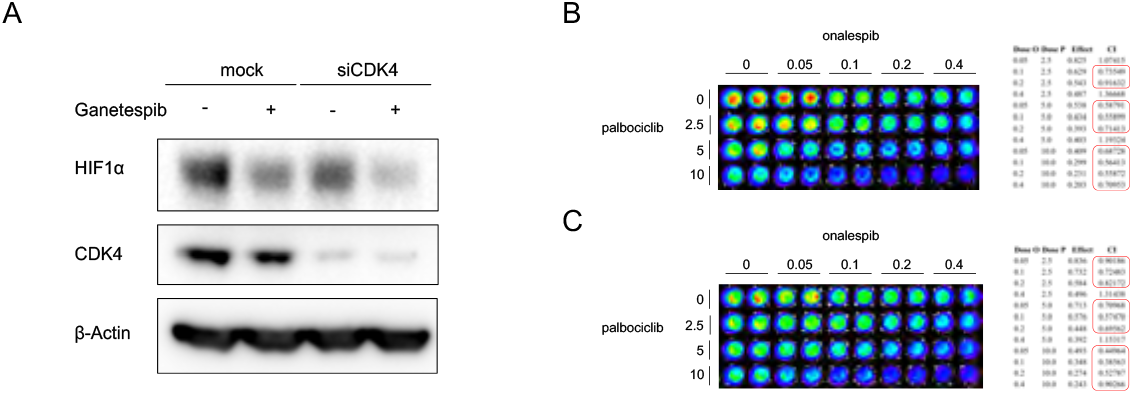
Dual inhibition of CDK4 and HSP90 decreases HIF1α level and synergistically inhibits cell viability in SW480 cells. (A) SW480 cells were treated with DMSO or ganetespib (1 μM) after 48 hours of knockdown of CDK4. (B, C) SW480 cells were treated with palbociclib and onalespib at the indicated doses for 72 hours in (B) normoxia and (C) hypoxia (0.5% O_2_).

**Supplementary figure 2.**
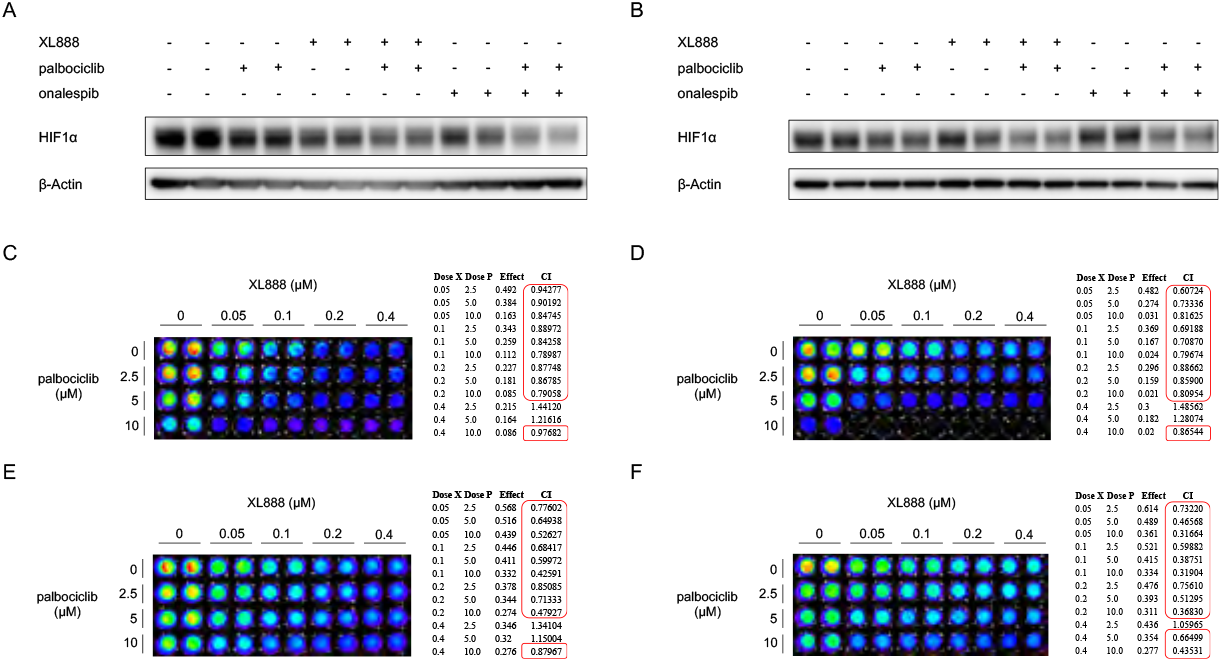
Combination treatment with the HSP90 inhibitor XL-888 and CDK4/6 inhibitor palbociclib inhibits HIF1α and cell viability in colorectal cancer. (A) HCT116 and (B) HT29 colon cancer cells were treated with indicated inhibitors (XL-888 at 0.05 μM, onalespib at 0.05 μM, and palbociclib at 10 μM) for 6 hours under hypoxia (0.5% O_2_). (C, D) In HCT116 and (E, F) SW480 colon cancer cells, XL-888 and palbociclib synergistically inhibit cell viability under (C, E) normoxia and (D, F) hypoxia (0.5% O_2_).

**Supplementary figure 3.**
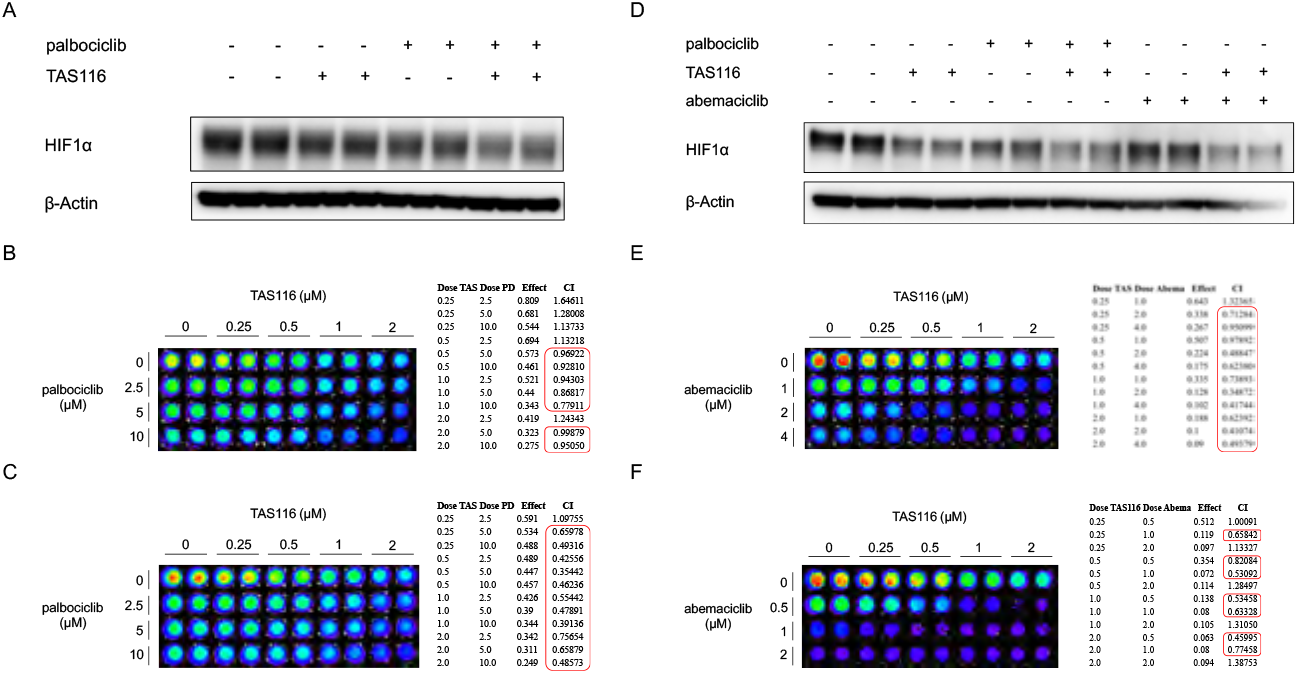
The combinatorial effect of CDK4/6 inhibitors with HSP90 inhibitors applies to alternative inhibitors in SW480 cells. (A) SW480 cells were treated with the indicated inhibitors (TAS-116 at 0.5 μM and palbociclib at 10 μM) for 6 hours under hypoxia (0.5% O_2_). (B, C) CDK4 inhibitor palbociclib and HSP90 inhibitor TAS-116 synergistically inhibit the viability of SW480 cells in (B) normoxia and (C) hypoxia (0.5% O_2_). (D) SW480 cells were treated with indicated inhibitors (TAS-116 at 0.5 μM and abemaciclib at 10 μM) for 6 hours under hypoxia (0.5% O_2_). (E, F) CDK4 inhibitor abemaciclib and HSP90 inhibitor TAS-116 synergistically inhibit the viability of SW480 cells in (B) normoxia and (C) hypoxia (0.5% O_2_).

**Supplementary figure 4.**
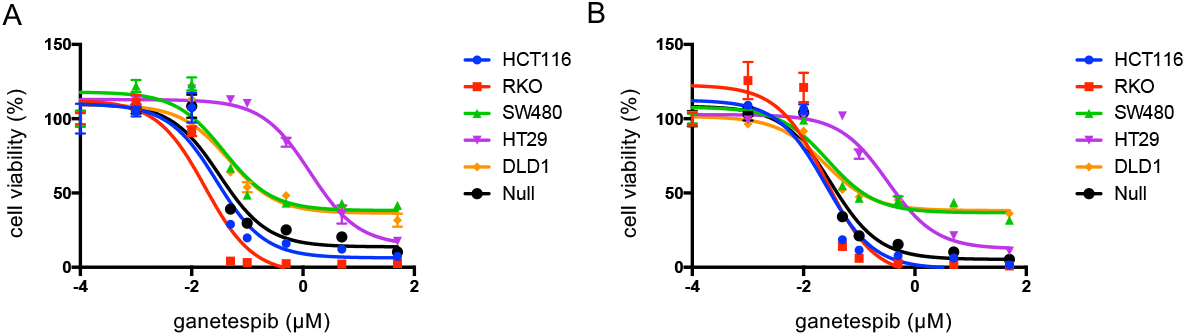
Dose response curve of ganetespib in colorectal cancer cell lines. (A) In normoxia or (B) in hypoxia (0.5% O_2_), cells were treated with increasing doses of ganetespib for 72 hours. Null: HCT116 p53^−/−^ colorectal cancer cells.

**Supplementary figure 5.**
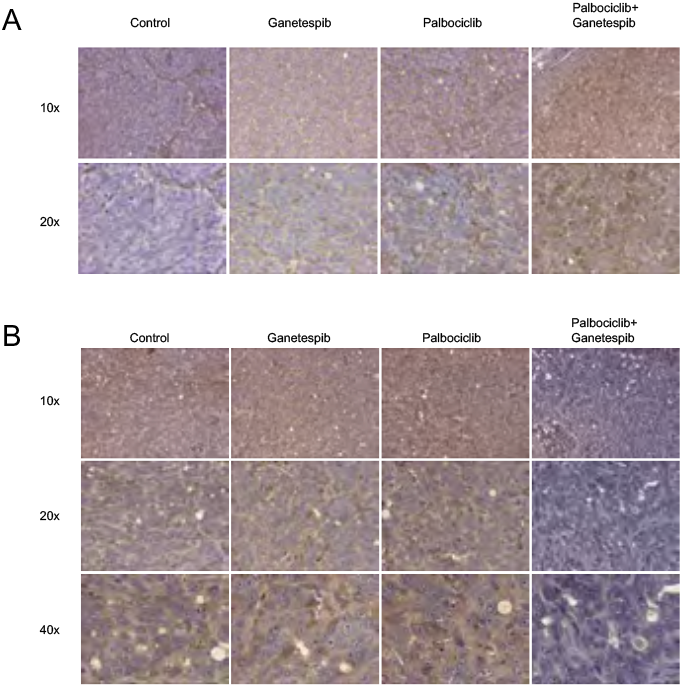
Combinatorial treatment by CDK4/6 inhibitor palbociclib and HSP90 inhibitor ganetespib increased caspase 3 cleavage and inhibited VEGF expression in xenograft tumors. (A) IHC staining of cleaved caspase 3. (B) IHC staining of VEGF.

**Supplementary figure 6.**
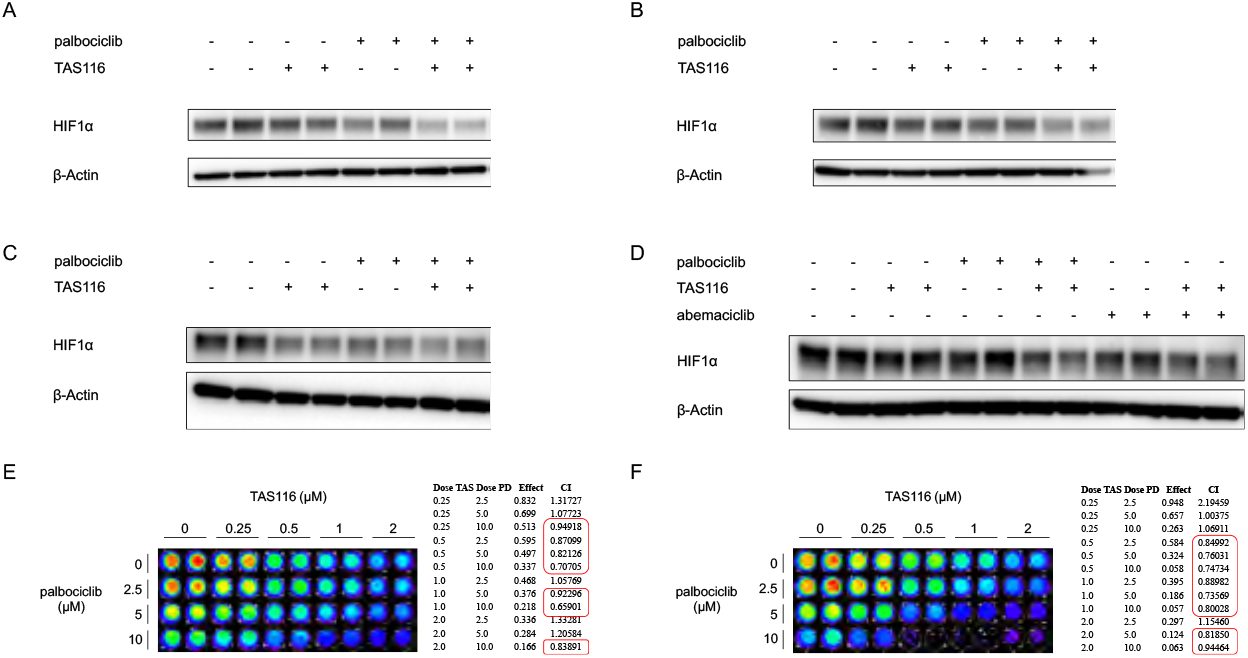
Dual inhibition of CDK4 and HSP90 inhibits HIF1α and cell viability in multiple cancer types. (A) ASPC1 and (B) HPAFII pancreatic cancer cells as well as (C) SKBR3 and (D) MDA-MB-361 breast cancer cells were treated with the indicated inhibitors (TAS-116 at 0.5 μM, palbociclib at 10 μM and abemaciclib at 1μM) for 6 hours under hypoxia (0.5% O_2_). (E) In normoxia and (F) hypoxia (0.5% O_2_), CDK4 inhibitor palbociclib and HSP90 inhibitor TAS-116 inhibit the viability of SKBR3 cells.

**Supplementary figure 7.**
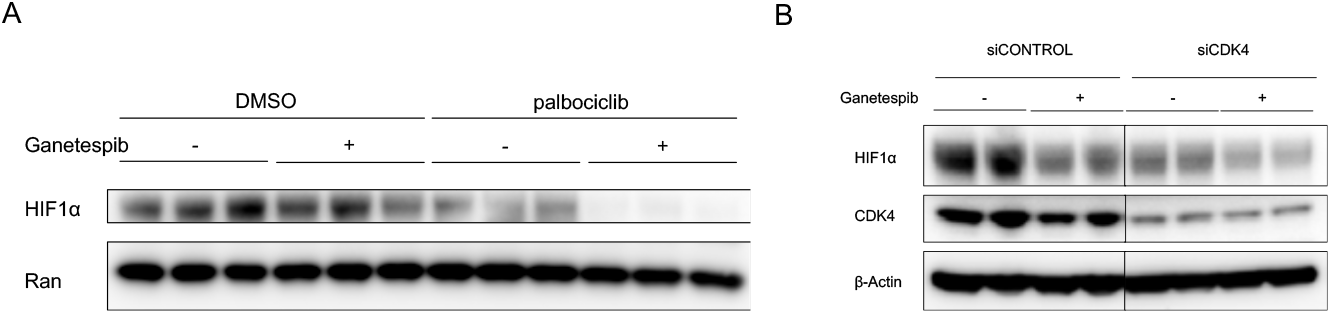
Dual inhibition of CDK4/6 and HSP90 robustly decreases the levels of HIF1α in multiple cancer cell types. (A) T98G cells were treated with palbociclib or ganetespib or the combination of both in hypoxia (0.5% O_2_) for 6 hours. (B) PC3 cells were treated with ganetespib for 6 hours under hypoxia (0.5% O_2_) after 48 hours of knockdown of CDK4.

**Supplementary figure 8.**
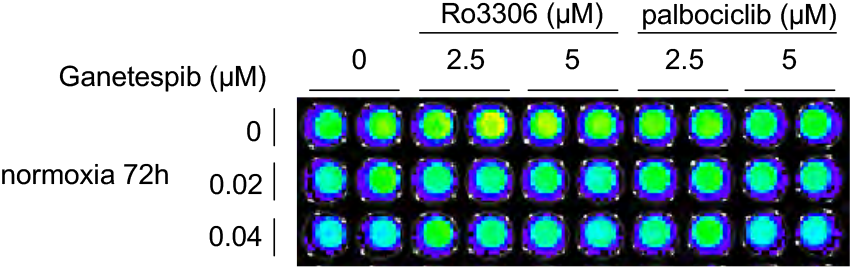
Combination of CDK1 inhibitor Ro-3306 or CDK4/6 inhibitor palbociclib and HSP90 inhibitor ganetespib does not induce cell death in WI38 normal cells in normoxia.

**Supplementary figure 9.**
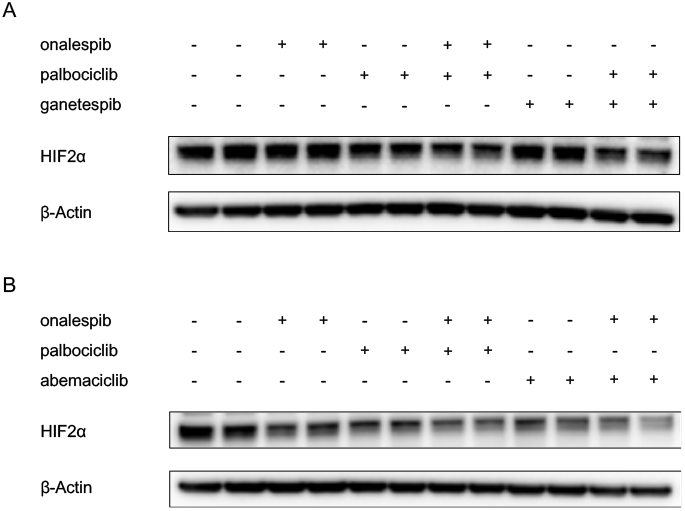
Dual inhibition of CDK4/6 and HSP90 slightly inhibits HIF2α in HCT116 cells. Cells were treated with the indicated inhibitors (ganetespib at 0.05 μM, onalespib at 0.05 μM, palbociclib and abemaciclib at 10 μM) for 6 hours under hypoxia (0.5% O_2_).

**Supplementary figure 10.**
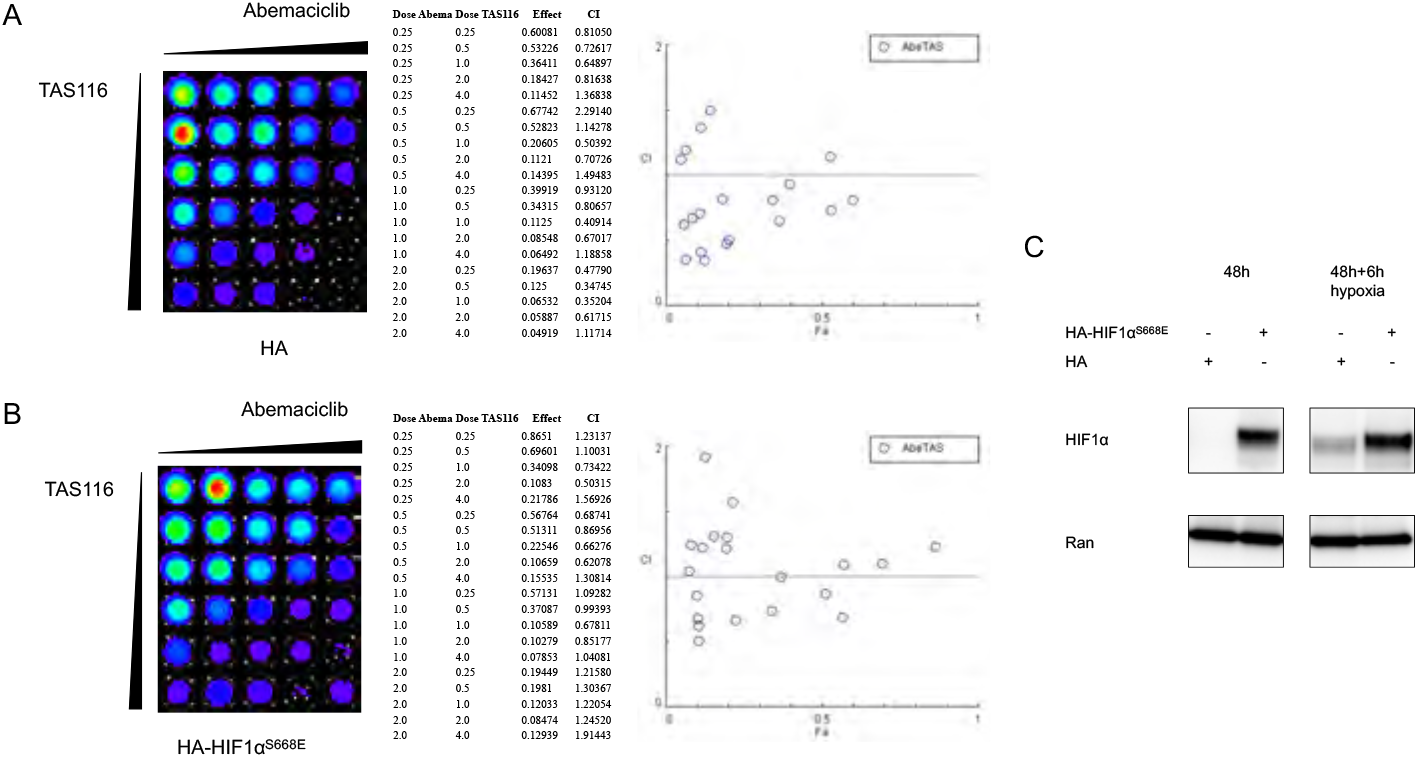
Overexpression of HIF1α^668E^ partially rescued the cell viability inhibition by combination of CDK4/6 and HSP90 inhibitor treatment under hypoxia. Cells were transfected with pcDNA3 plasmid carrying (A) HA tag or (B) HA-HIF1α^668E^ for 48 hours, and subsequently treated with the indicated drug combinations for 72 hours under hypoxia (0.5% O_2_). (C) HIF1α overexpression by HA-HIF1α^668E^ at 48 hours post transfection or 48 hours transfection plus 6 hours 0.5% O_2_ hypoxia treatment.

**Supplementary figure 11.**
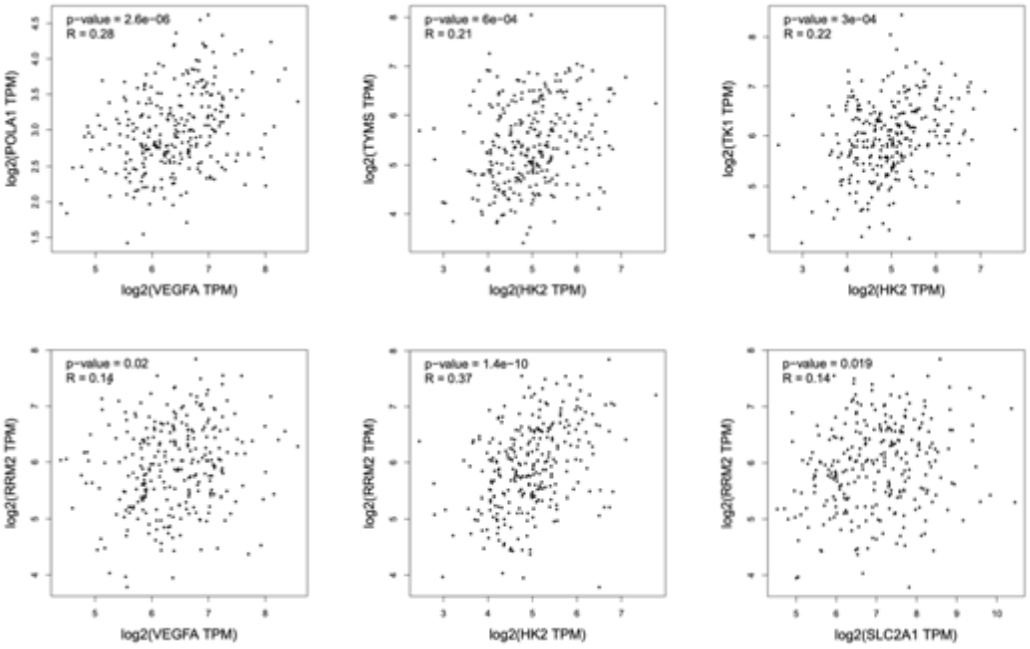
Pearson correlation analysis between HIF1α and E2F target genes in colon adenocarcinoma. The analysis is performed using the GEPIA online tool based on TCGA colon adenocarcinoma data.

**Supplementary figure 12.**
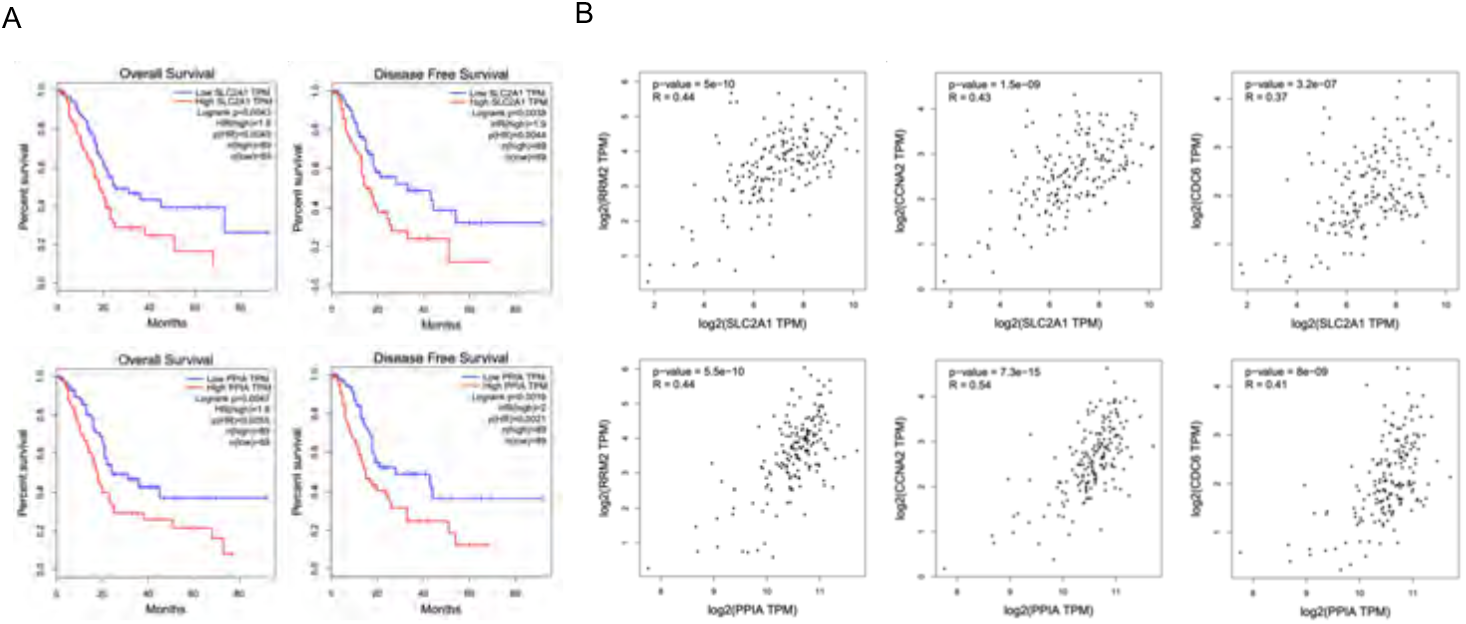
Analysis of TCGA data in pancreatic adenocarcinoma (PAAD). (A) Overexpression of HIF1α target genes SLC2A1 and PPIA is correlated with poor overall survival and disease-free survival in PAAD patients. (B) Correlations between HIF1α target genes (SLC2A1 and PPIA) and E2F target genes (RRM2, CCNA2 and CDC6) based on TCGA PAAD data.

## Reference

1 Semenza, G. L. Hypoxia-inducible factors in physiology and medicine. Cell 148, 399–408, doi:10.1016/j.cell.2012.01.021 (2012).

2 Carmeliet, P. & Jain, R. K. Angiogenesis in cancer and other diseases. Nature 407, 249–257, doi:10.1038/35025220 (2000).

3 Wilson, W. R. & Hay, M. P. Targeting hypoxia in cancer therapy. Nature reviews. Cancer 11, 393–410, doi:10.1038/nrc3064 (2011).

4 Minassian, L. M., Cotechini, T., Huitema, E. & Graham, C. H. Hypoxia-Induced Resistance to Chemotherapy in Cancer. Adv Exp Med Biol 1136, 123–139, doi:10.1007/978-3-030-12734-3_9 (2019).

5 Horsman, M. R. & Overgaard, J. The impact of hypoxia and its modification of the outcome of radiotherapy. J Radiat Res 57 1, i90–i98, doi:10.1093/jrr/rrw007 (2016).

6 Yun, Z. & Lin, Q. Hypoxia and regulation of cancer cell stemness. Adv Exp Med Biol 772, 41–53, doi:10.1007/978-1-4614-5915-6_2 (2014).

7 Nobre, A. R., Entenberg, D., Wang, Y., Condeelis, J. & Aguirre-Ghiso, J. A. The Different Routes to Metastasis via Hypoxia-Regulated Programs. Trends Cell Biol 28, 941–956, doi:10.1016/j.tcb.2018.06.008 (2018).

8 Masson, N. & Ratcliffe, P. J. Hypoxia signaling pathways in cancer metabolism: the importance of co-selecting interconnected physiological pathways. Cancer Metab 2, 3, doi:10.1186/2049-3002-2-3 (2014).

9 Kaelin, W. G., Jr. Cancer and altered metabolism: potential importance of hypoxia-inducible factor and 2-oxoglutarate-dependent dioxygenases. Cold Spring Harb Symp Quant Biol 76, 335–345, doi:10.1101/sqb.2011.76.010975 (2011).

10 Chen, S. & Sang, N. Hypoxia-Inducible Factor-1: A Critical Player in the Survival Strategy of Stressed Cells. J Cell Biochem 117, 267–278, doi:10.1002/jcb.25283 (2016).

11 Baba, Y. et al. HIF1A overexpression is associated with poor prognosis in a cohort of 731 colorectal cancers. The American journal of pathology 176, 2292–2301, doi:10.2353/ajpath.2010.090972 (2010).

12 Dekervel, J. et al. Hypoxia-driven gene expression is an independent prognostic factor in stage II and III colon cancer patients. Clinical cancer research: an official journal of the American Association for Cancer Research 20, 2159–2168, doi:10.1158/1078-0432.ccr-13-2958 (2014).

13 Nishimoto, A. et al. HIF-1alpha activation under glucose deprivation plays a central role in the acquisition of anti-apoptosis in human colon cancer cells. International journal of oncology 44, 2077–2084, doi:10.3892/ijo.2014.2367 (2014).

14 Zhang, W. et al. HIF-1alpha Promotes Epithelial-Mesenchymal Transition and Metastasis through Direct Regulation of ZEB1 in Colorectal Cancer. PloS one 10, e0129603, doi:10.1371/journal.pone.0129603 (2015).

15 Forsythe, J. A. et al. Activation of vascular endothelial growth factor gene transcription by hypoxia-inducible factor 1. Molecular and cellular biology 16, 4604–4613 (1996).

16 Ivan, M. et al. HIFalpha targeted for VHL-mediated destruction by proline hydroxylation: implications for O2 sensing. Science (New York, N.Y) 292, 464–468, doi:10.1126/science.1059817 (2001).

17 Kaelin, W. G., Jr. The VHL Tumor Suppressor Gene: Insights into Oxygen Sensing and Cancer. Trans Am Clin Climatol Assoc 128, 298–307 (2017).

18 Cowey, C. L. & Rathmell, W. K. VHL gene mutations in renal cell carcinoma: role as a biomarker of disease outcome and drug efficacy. Current oncology reports 11, 94–101 (2009).

19 Peng, X.-H. et al. Cross-talk between Epidermal Growth Factor Receptor and Hypoxia-inducible Factor-1α Signal Pathways Increases Resistance to Apoptosis by Up-regulating Survivin Gene Expression. Journal of Biological Chemistry 281, 25903–25914, doi:10.1074/jbc.M603414200 (2006).

20 Talks, K. L. et al. The expression and distribution of the hypoxia-inducible factors HIF-1alpha and HIF-2alpha in normal human tissues, cancers, and tumor-associated macrophages. The American journal of pathology 157, 411–421 (2000).

21 Dang, Eric V. et al. Control of TH17/Treg Balance by Hypoxia-inducible Factor 1. Cell 146, 772–784, doi:10.1016/j.cell.2011.07.033.

22 Mayes, P. A. et al. Overcoming hypoxia-induced apoptotic resistance through combinatorial inhibition of GSK-3beta and CDK1. Cancer Res 71, 5265–5275, doi:10.1158/0008-5472.CAN-11-1383 (2011).

23 Warfel, N. A., Dolloff, N. G., Dicker, D. T., Malysz, J. & El-Deiry, W. S. CDK1 stabilizes HIF-1alpha via direct phosphorylation of Ser668 to promote tumor growth. Cell cycle (Georgetown, Tex.) 12, 3689–3701, doi:10.4161/cc.26930 (2013).

24 Isaacs, J. S. et al. Hsp90 regulates a von Hippel Lindau-independent hypoxia-inducible factor-1 alpha-degradative pathway. The Journal of biological chemistry 277, 29936–29944, doi:10.1074/jbc.M204733200 (2002).

25 Gradin, K. et al. Functional interference between hypoxia and dioxin signal transduction pathways: competition for recruitment of the Arnt transcription factor. Molecular and cellular biology 16, 5221–5231 (1996).

26 Huang, T. et al. Expression of Hsp90alpha and cyclin B1 were related to prognosis of esophageal squamous cell carcinoma and keratin pearl formation. International journal of clinical and experimental pathology 7, 1544–1552 (2014).

27 McCarthy, M. M. et al. HSP90 as a marker of progression in melanoma. Annals of oncology: official journal of the European Society for Medical Oncology 19, 590–594, doi:10.1093/annonc/mdm545 (2008).

28 Tian, W. L. et al. High expression of heat shock protein 90 alpha and its significance in human acute leukemia cells. Gene 542, 122–128, doi:10.1016/j.gene.2014.03.046 (2014).

29 Asghar, U., Witkiewicz, A. K., Turner, N. C. & Knudsen, E. S. The history and future of targeting cyclin-dependent kinases in cancer therapy. Nature reviews. Drug discovery 14, 130–146, doi:10.1038/nrd4504 (2015).

30 Proia, D. A. & Bates, R. C. in Heat Shock Protein-Based Therapies (eds Alexzander A. A. Asea, Naif N. Almasoud, Sunil Krishnan, & Punit Kaur) 289–322 (Springer International Publishing, 2015).

31 Vassilev, L. T. et al. Selective small-molecule inhibitor reveals critical mitotic functions of human CDK1. Proceedings of the National Academy of Sciences of the United States of America 103, 10660–10665, doi:10.1073/pnas.0600447103 (2006).

32 Turner, N. C. et al. Palbociclib in Hormone-Receptor-Positive Advanced Breast Cancer. The New England journal of medicine 373, 209–219, doi:10.1056/NEJMoa1505270 (2015).

33 Vijayaraghavan, S. et al. CDK4/6 and autophagy inhibitors synergistically induce senescence in Rb positive cytoplasmic cyclin E negative cancers. Nature communications 8, 15916, doi:10.1038/ncomms15916 (2017).

34 Ying, W. et al. Ganetespib, a unique triazolone-containing Hsp90 inhibitor, exhibits potent antitumor activity and a superior safety profile for cancer therapy. Molecular cancer therapeutics 11, 475–484, doi:10.1158/1535-7163.mct-11-0755 (2012).

35 Nagaraju, G. P. et al. Antiangiogenic effects of ganetespib in colorectal cancer mediated through inhibition of HIF-1alpha and STAT-3. Angiogenesis 16, 903–917, doi:10.1007/s10456-013-9364-7 (2013).

36 He, S. et al. The HSP90 inhibitor ganetespib has chemosensitizer and radiosensitizer activity in colorectal cancer. Investigational new drugs 32, 577–586, doi:10.1007/s10637-014-0095-4 (2014).

37 Takayama, T., Miyanishi, K., Hayashi, T., Sato, Y. & Niitsu, Y. Colorectal cancer: genetics of development and metastasis. Journal of gastroenterology 41, 185–192, doi:10.1007/s00535-006-1801-6 (2006).

38 Cui, Y. An Integrative Procedure for Apoptosis Identification and Measurement. (2006).

39 Kaufmann, S. H., Desnoyers, S., Ottaviano, Y., Davidson, N. E. & Poirier, G. G. Specific proteolytic cleavage of poly(ADP-ribose) polymerase: an early marker of chemotherapy-induced apoptosis. Cancer research 53, 3976–3985 (1993).

40 Franken, N. A. P., Rodermond, H. M., Stap, J., Haveman, J. & van Bree, C. Clonogenic assay of cells in vitro. Nature Protocols 1, 2315, doi:10.1038/nprot.2006.339 (2006).

41 Nagaraju, G. P., Bramhachari, P. V., Raghu, G. & El-Rayes, B. F. Hypoxia inducible factor-1alpha: Its role in colorectal carcinogenesis and metastasis. Cancer letters 366, 11–18, doi:10.1016/j.canlet.2015.06.005 (2015).

42 Liang, C.-C., Park, A. Y. & Guan, J.-L. In vitro scratch assay: a convenient and inexpensive method for analysis of cell migration in vitro. Nature Protocols 2, 329, doi:10.1038/nprot.2007.30 (2007).

43 Chen, X. et al. XBP1 promotes triple-negative breast cancer by controlling the HIF1alpha pathway. Nature 508, 103–107, doi:10.1038/nature13119 (2014).

44 Liu, J. et al. Parkin targets HIF-1alpha for ubiquitination and degradation to inhibit breast tumor progression. Nature communications 8, 1823, doi:10.1038/s41467-017-01947-w (2017).

45 Shukla, S. K. et al. MUC1 and HIF-1alpha Signaling Crosstalk Induces Anabolic Glucose Metabolism to Impart Gemcitabine Resistance to Pancreatic Cancer. Cancer cell 32, 71–87.e77, doi:10.1016/j.ccell.2017.06.004 (2017).

46 Palazon, A. et al. An HIF-1alpha/VEGF-A Axis in Cytotoxic T Cells Regulates Tumor Progression. Cancer cell 32, 669–683.e665, doi:10.1016/j.ccell.2017.10.003 (2017).

47 Krzywinska, E. et al. Loss of HIF-1alpha in natural killer cells inhibits tumour growth by stimulating non-productive angiogenesis. Nature communications 8, 1597, doi:10.1038/s41467-017-01599-w (2017).

48 Noman, M. Z. et al. PD-L1 is a novel direct target of HIF-1alpha, and its blockade under hypoxia enhanced MDSC-mediated T cell activation. The Journal of experimental medicine 211, 781–790, doi:10.1084/jem.20131916 (2014).

49 Spencer, J. A. et al. Direct measurement of local oxygen concentration in the bone marrow of live animals. Nature 508, 269–273, doi:10.1038/nature13034 (2014).

50 Giambra, V. et al. Leukemia stem cells in T-ALL require active Hif1alpha and Wnt signaling. Blood 125, 3917–3927, doi:10.1182/blood-2014-10-609370 (2015).

51 Zou, J. et al. Notch1 is required for hypoxia-induced proliferation, invasion and chemoresistance of T-cell acute lymphoblastic leukemia cells. Journal of hematology & oncology 6, 3, doi:10.1186/1756-8722-6-3 (2013).

52 Fry, D. W. et al. Specific inhibition of cyclin-dependent kinase 4/6 by PD 0332991 and associated antitumor activity in human tumor xenografts. Molecular cancer therapeutics 3, 1427–1438 (2004).

53 Rosner, M., Pham, H. T. T., Moriggl, R. & Hengstschlager, M. Human stem cells alter the invasive properties of somatic cells via paracrine activation of mTORC1. Nature communications 8, 595, doi:10.1038/s41467-017-00661-x (2017).

54 Kitajima, S. et al. Hypoxia-inducible factor-1alpha promotes cell survival during ammonia stress response in ovarian cancer stem-like cells. Oncotarget 8, 114481–114494, doi:10.18632/oncotarget.23010 (2017).

55 Conley, S. J. et al. Antiangiogenic agents increase breast cancer stem cells via the generation of tumor hypoxia. Proceedings of the National Academy of Sciences of the United States of America 109, 2784–2789, doi:10.1073/pnas.1018866109 (2012).

56 Liao, D., Corle, C., Seagroves, T. N. & Johnson, R. S. Hypoxia-inducible factor-1alpha is a key regulator of metastasis in a transgenic model of cancer initiation and progression. Cancer research 67, 563–572, doi:10.1158/0008-5472.can-06-2701 (2007).

57 Singleton, D. C. et al. Hypoxic regulation of RIOK3 is a major mechanism for cancer cell invasion and metastasis. Oncogene 34, 4713–4722, doi:10.1038/onc.2014.396 (2015).

58 Gilkes, D. M. et al. Collagen prolyl hydroxylases are essential for breast cancer metastasis. Cancer research 73, 3285–3296, doi:10.1158/0008-5472.can-12-3963 (2013).

59 Corzo, C. A. et al. HIF-1alpha regulates function and differentiation of myeloid-derived suppressor cells in the tumor microenvironment. The Journal of experimental medicine 207, 2439–2453, doi:10.1084/jem.20100587 (2010).

60 Doedens, A. L. et al. Macrophage expression of hypoxia-inducible factor-1 alpha suppresses T-cell function and promotes tumor progression. Cancer research 70, 7465–7475, doi:10.1158/0008-5472.can-10-1439 (2010).

61 Clambey, E. T. et al. Hypoxia-inducible factor-1 alpha-dependent induction of FoxP3 drives regulatory T-cell abundance and function during inflammatory hypoxia of the mucosa. Proceedings of the National Academy of Sciences of the United States of America 109, E2784–2793, doi:10.1073/pnas.1202366109 (2012).

62 Barsoum, I. B., et al. A mechanism of hypoxia-mediated escape from adaptive immunity in cancer cells. Cancer research 74, 665–674, doi:10.1158/0008-5472.can-13-0992 (2014).

63 Goel, S. et al. “CDK4/6 inhibition triggers anti-tumour immunity.” Nature 548, 471–475. doi:10.1038/nature23465 (2017).

64 Deng, J. et al. “CDK4/6 Inhibition Augments Antitumor Immunity by Enhancing T-cell Activation.” Cancer discovery 8, 216–233. doi:10.1158/2159-8290.CD-17-0915 (2018).

65 Zhang, J. et al. “Cyclin D-CDK4 kinase destabilizes PD-L1 via cullin 3-SPOP to control cancer immune surveillance.” Nature 553, 91–95. doi:10.1038/nature25015 (2018).

66 Kondo, K., et al. Inhibition of HIF is necessary for tumor suppression by the von Hippel-Lindau protein. Cancer cell 1, 237–246, doi:10.1016/s1535-6108(02)00043-0 (2002).

